# When spatial attention cannot be divided: Quadrantic enhancement of early visual processing across task–relevant and irrelevant locations

**DOI:** 10.1101/2024.03.19.585785

**Authors:** Mert Özkan, Viola Störmer

## Abstract

Spatial attention enables us to select regions of space and prioritize visual processing at the attended locations. Previous research has shown that spatial attention can be flexibly tuned to broader or narrower regions in space, and in some cases be split amongst multiple locations. Here, we investigate how attentional resources are distributed within a visual quadrant when participants are instructed to either focus attention narrowly, broadly, or split attention among two non-contiguous locations. Using a combination of behavior and steady-state visual-evoked potentials (SSVEP), the oscillatory response of the visual cortex to incoming flickering stimuli, we find clear evidence for ineffective splitting of spatial attention within a visual quadrant. Importantly, by assessing visual-cortical processing across locations at a high spatial resolution (by flickering nearby locations at distinct frequencies), our results reveal that attention was distributed in the exact same manner regardless of whether participants were instructed to attend broadly across a large region of space, or divide attention amongst two non-contiguous locations: In both cases, the intermediate location showed the strongest boost in visual-cortical processing, no matter whether it was the center of the attended region (broad-focus condition), or the uncued, to-be-ignored location (split-focus condition). Thus, the present study provides strong evidence that when trying to attend to multiple separate locations within a visual quadrant, sustained attention inadvertently enhances visual processing at the intermediate location even when it is detrimental to task performance.

## Introduction

Spatial attention allows us to select specific regions of the visual field and prioritize processing of information at these attended locations. This prioritization results in increased behavioral performance for stimuli falling within the selected region relative to other stimuli, as well as changes in neural processing in the early visual cortex (Reynolds & Chelazzi, 2004), for example through increases in neural gain (Hillyard et al., 1998; Itthipuripat et al., 2014), sharper tuning of the neural responses (Anton-Erxleben et al., 2009), or noise reduction (Lu & Dosher, 2004). Early models often conceptualized spatial attention as a “spotlight” that highlights the relevant region of space (Desimone & Duncan, 1995), and proposed that the prioritized region could flexibly vary in size, as if attention was “zooming” in and out, depending on task demands (e.g. the zoom-lens model, Eriksen & St James, 1986). Consistent with this, recent models – for example, the normalization model of attention (Reynolds & Heeger, 2009) – put a strong emphasis on the size of the attentional focus, suggesting that several aspects of behavioral performance can be explained by a combination of the breadth of the attentional focus as well as the size of the stimulus in a given task (Herrmann et al., 2010). Thus, it is commonly accepted that spatial attention can be focused narrowly on a small region in space or more broadly across larger parts of the visual field, depending on task demands.

A related question that has been long debated in the literature on spatial attention is whether resources can be split across noncontiguous regions of space. While several studies found evidence for the existence of multiple attentional foci (Awh & Pashler, 2000; McMains & Somers, 2005; Morawetz et al., 2007; Müller et al., 2003; Niebergall et al., 2011; Störmer et al., 2013), other studies yielded less clear results as to when split–foci can arise (Alvarez et al., 2012; for a review see Jans et al., 2010). One important factor to consider when evaluating whether attention can be split across distinct regions in space is the cortical architecture of the visual system, in particular the anatomical division in the early visual cortex where the left visual cortex processes the right visual field and vice versa. These hemifield boundaries strongly constrain how attention can be split: attending to multiple target locations across the visual hemifields comes at little to no cost, whereas attending to multiple locations within a hemifield results in a reduction in performance (Alvarez et al., 2012; Alvarez & Cavanagh, 2005; Awh & Pashler, 2000; Bichot et al., 1999; Störmer et al., 2014). For example, in a seminal study, Awh and Pashler (2000) used a divided spatial attention task to test visual discrimination performance at cued locations as well as an uncued location that was located in between the two attended locations, and found that performance at the intervening uncued location was not significantly different from at least one cued location when stimuli were presented within a hemifield; providing behavioral support for the notion that splitting attention across two non-contiguous regions within a visual half-field is inefficient.

Other evidence comes from a series of electroencephalography (EEG) studies that used steady-state visually evoked potentials (SSVEPs) to directly measure early visual-cortical processing of attended and unattended locations. The SSVEP is the oscillatory response generated in the visual cortex, evoked by and phase-locked to a flickering visual stimulus (Regan, 1988). Importantly, attending to a specific stimulus directly modulates the response amplitude of the given stimulus frequency as compared to unattended stimuli (Andersen et al., 2015; Morgan et al., 1996, Müller et al., 2003). In a series of studies, Walter and colleagues (2014; 2016) tested differences between attending within– versus across–hemifield locations and found that the across–hemifield attention condition reliably induced modulations at attended locations whereas the attentional modulations within-hemifield condition were weaker or unreliable (see also, Malinowski et al., 2007). Similarly, another study that examined attentional modulations of target– evoked SSVEPs relative to distractor–evoked SSVEPs within a hemifield or visual quadrant failed to observe reliable attentional modulation of target-evoked activity (Itthipuripat et al., 2013). Finally, data from a multiple–object tracking task where participants were asked to keep track of multiple moving targets within or across hemifields showed reliable attentional modulations only during across-hemifield tracking but not within-hemifield tracking (Störmer et al., 2014). Collectively, these studies indicate that early attentional selection of multiple locations is compromised when these locations fall within a single visual half-field or quadrant.

Interestingly, none of these existing studies have systematically tested why attending to multiple locations within a hemifield is less efficient, or in other words, how attention is distributed when participants are instructed to attend to two noncontiguous regions in a single hemifield or visual quadrant. Given that many studies failed to find any reliable attentional modulations in the within–hemifield conditions, it is particularly difficult to interpret these null results. For example, it is unclear whether cueing participants to attend to multiple locations within a hemifield results in broadly–tuned attention across cued and uncued locations, or a complete failure of attentional enhancement.

To directly assess how spatial attention operates within a hemifield, we here compared three cueing conditions (see Figure 1B): (1) a split–focus condition where participants were cued to attend to two noncontiguous locations with an intermediate, uncued location; (2) a broad –focus condition where participants were cued to attend to a large contiguous region that included the same intermediate location; (3) a narrow–focus condition where participants were cued to attend to a small region, which served as a control. The main question of interest was whether any attention effects would differ between the split– and broad–focus conditions in terms of behavioral performance (Experiment 1) or early visual-cortical responses (as indexed by SSVEPs, Experiments 2 & 3). We hypothesized that if cueing participants to attend to two noncontiguous locations results in the same distribution of resources as being cued to attend broadly to the entire region, there should be no difference between the multifocal and broad–focus conditions. However, if cueing participants to attend to two locations results in less efficient attentional enhancement overall, or attending to only one of the two cued locations, this should result in less or no modulation on average, relative to attending broadly. To preview our results, we observed reliable cueing effects (Experiment 1-3) and attentional modulations in early visual processing (Experiments 2-3) across all experiments; however, the intermediate location was processed similarly regardless of whether participants were instructed to ignore it (split–focus condition) or to attend to it (broad–focus condition). These findings suggest that even if the task demands splitting the focus of attention within a visual quadrant, spatial attention acts to enhance all locations, very similar to when the task requires a broadening of the focus of attention.

**Figure 1.**
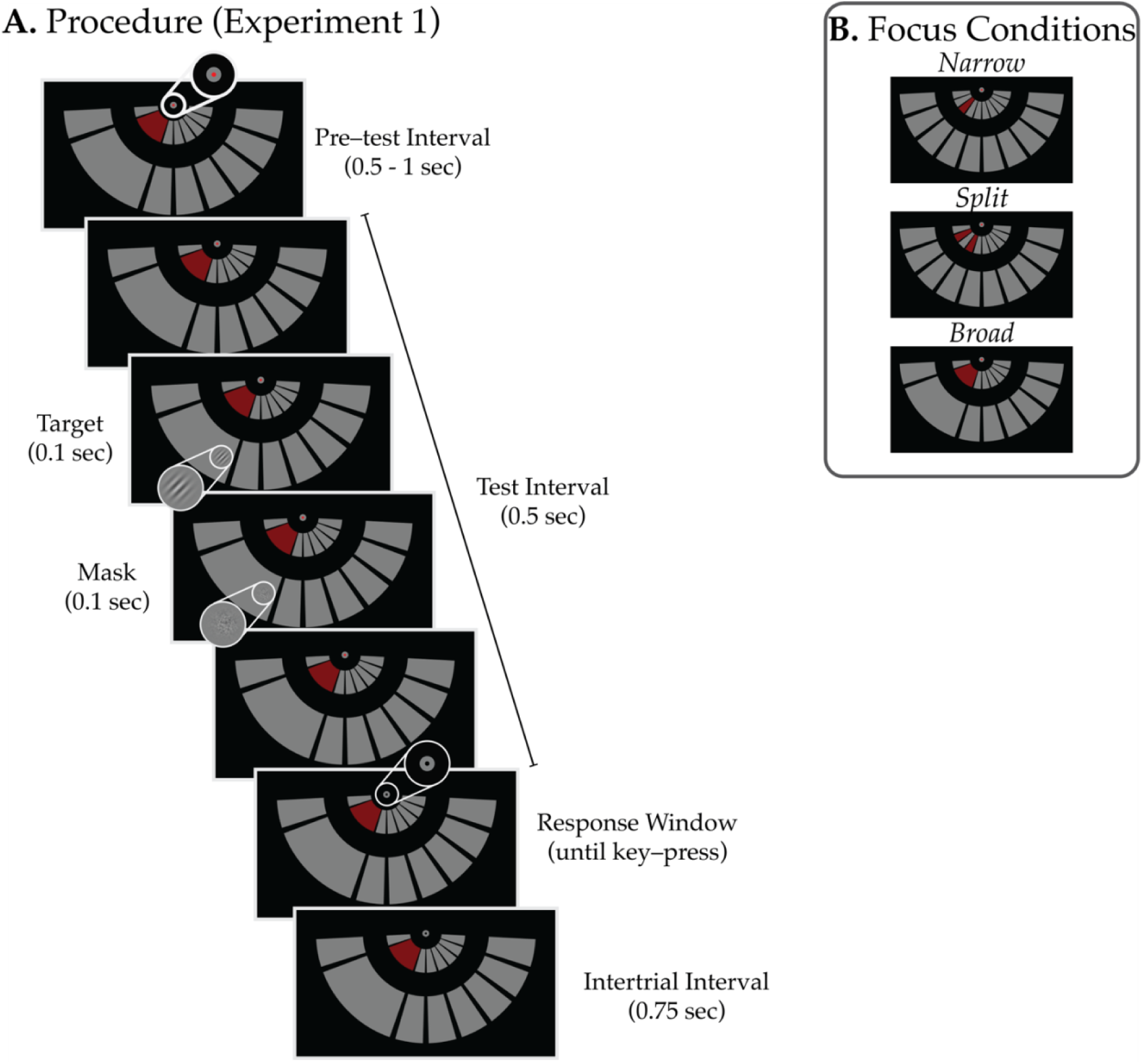
(A) *Procedure for Experiment* î. Each trial started with a pre-test interval (0.5 -1 sec), indicated by the fixation color change, which was followed by the test interval where the target stimulus appeared for 0.1 sec and was immediately succeeded by a white noise mask (0.1 sec), at a random time. The fixation color turned to black indicating the response window and the participant reported the orientation (+ or - 45° from vertical) by keypress. An intertrĩal interval of 0.75 sec followed the response. (B) *Focus Conditions.* Participants either attended to a narrow location (top), two non-contiguous locations (split, middle), or a broad location (bottom) indicated by the red inner arcs that served at spatial attention cues. 75% of the targets appeared on the cued location(s) and 25% of the targets appeared on uncued locations that were immedi­ately adjacent to a cued location. Focus conditions were counterbalanced across the left and right lower visual field.

## Experiment I

In the first experiment, we tested how effectively participants can attend to two noncontiguous regions relative to attending broadly to a larger region in space using behavioral methods. Participants performed a probabilistic cueing task where they were instructed to attend to a single small location (narrow–focus), one relatively large location (broad–focus), or two noncontiguous locations with an intermediate uncued region (split–focus) within a visual quadrant (see Figure 1B). Critically, we tested visual discrimination sensitivity both in the cued and the uncued locations. If attention can be divided effectively, participants’ performance should differ at the intermediate region depending on whether it was more likely to contain a target (cued in the broad–focus condition) or not (uncued in the split–focus condition).

### Methods

#### Participants

30 undergraduate students from Dartmouth College, aged between 18 and 35 years, participated in the experiment. All participants had normal or corrected-to-normal vision. Prior to the experiment, participants signed an informed consent form as approved by the Institutional Review Board at Dartmouth College. The study protocol was approved by the Committee for the Protection of Human Subjects.

#### Apparatus

The experiment was programmed in MATLAB R2020b using the PsychToolbox-3 (Brainard, 1997; Pelli, 1997). Testing was completed on a PC running Windows 10. The stimuli were displayed in an otherwise dark room on an ASUS VG279QM monitor (1920 × 1800 pixels at 239.7 Hz) placed 81 cm from the participant. The participant’s head was stabilized by a chin and forehead rest. Participants’ eyes were tracked online via a desktop–mount Eyelink 1000 Plus eye-tracker (SR Research) sampled at 500 Hz.

#### Stimuli

Figure 1 shows the stimulus configuration and trial sequence used in the experiment. Stimuli were presented on a black background. Participants were instructed to fixate on a central fixation dot comprising a small circle (0.1 degree-visual-angle (dva) radius) superimposed on a larger (0.25 dva) gray (RGB: [127, 127, 127]) circle. The inner circle changed color to indicate the trial epochs. The stimulus display was composed of two arcs that were partitioned into smaller segments, one arc nearby the central fixation circle, and the other one larger and further away from fixation; the former served to cue participants to specific locations on the larger arc; the latter one contained the locations the participant was attending. The partitioned arcs were created by drawing 10 equally sized segments along a circle centered around the fixation point. The outer arc extended from 8 dva to 14 dva away from the center and spanned 14.7°; that is, if two vectors were drawn both starting from the fixation point and each ending on the opposite sides of the arc the degree between these two vectors would be 14.7°. The arcs were separated by 3° from one another and 1.5° away from the vertical and horizontal meridians. In the broad –focus blocks (see *Procedure*) the three central segments at the cued side were merged to create a large contiguous area (50.1°). A scaled version of the partitioned arc was placed between 2 and 5 dva away from the fixation as a cue to indicate which locations to attend. The cued locations were highlighted in red (RGBA: [255, 0, 0, 127]). A Gabor patch (2 dva in radius, 0.1 spatial frequency) with a ±45° tilt from vertical was presented with a Gaussian mask (σ = .5) as a test stimulus. The contrast of the patch was determined individually for each participant through a thresholding procedure (see *Procedure*). A white noise mask of 50% Michelson contrast immediately followed the test probes. The mask matched the test stimulus’ size and a Gaussian mask with the same parameters as the test stimulus’ was superimposed on it.

### Procedure

For each participant, the experiment followed the same structure: they started with a guided practice including instructions, followed by an uninterrupted practice, an adaptive thresholding procedure, and lastly the main experiment. During the instructions, participants performed the task (60 trials) with high–contrast test probes (50%) that were easily visible. Then, the practice block (60 trials) presented lower-contrast probes (20%), making the task more challenging, and the eyetracker was used to monitor participants’ eye gaze and ensure fixation. The subsequent adaptive thresholding procedure was implemented using the Quest algorithm (Watson & Pelli, 1983) to estimate the test stimulus’ contrast necessary for ∼75% performance for each participant. During thresholding, participants were cued to attend broadly to the whole left or right visual quadrant and all five locations within a quadrant were merged to create one large contiguous region. The test probes appeared in one of the 9 equidistant positions at equal probability. The cued side changed after every 6 trials. The algorithm was updated after every response regardless of probe location or side. The estimated contrast threshold was used during the main experimental blocks.

Each experimental block started with the cue display (inner arc) indicating the high– probability locations where 75% of the test stimuli would be presented. The remaining 25% of the probes appeared at one of the neighboring locations. Participants were informed about these probabilities and instructed to covertly attend to the cued locations, while keeping their gaze in the center of the display. Each trial started with the fixation dot turning red to indicate the pre–test interval (jittered between 0.5 - 1 s, uniform distribution) where no test probe was displayed. At a random time within the next 0.5 s (test interval) a test stimulus appeared for 0.1 s, immediately followed by a masking stimulus (0.1 s), with the constraint that the test stimulus needed to be displayed a minimum of 0.2 s before the response period to make sure both the Gabor patch and the mask would be shown. Then, the fixation color turned black to indicate the response window for participants. Participants were instructed to report the orientation (+ or - 45° tilted away from vertical) of the Gabor patch using the “down arrow” or “left arrow” keys, respectively. The trial ended after the forced choice response and a 0.75–second–long intertrial interval (ITI) followed prior to the start of the next trial.

We used three types of cue conditions where the cued locations, when combined, had 75% chance of containing a test probe and the uncued locations had 25% (see Figure 1B): (1) in the narrow–focus condition one of the three central locations was the high–probability (75%) region whereas the two locations that are adjacent to it had equal probability (12.5% each) of containing a test probe; (2) in the split–focus condition, the two cued locations were the high– probability regions with an equal chance of containing the test probe (37.5% each), and all three locations adjacent to these cued locations including the intermediate region had an equal probability of containing a test probe (8.33% each); (3) in the broad–focus condition, all three central locations were merged eliminating the boundaries between them, and the entire merged segment was cued. In order to make sure participants distributed their attentional resources over the entire cued region, the cued targets could appear at 5 equidistant locations instead of 3 so that each cued location had 15% chance of containing a target (75% in total) and the uncued test probes appeared in one of the uncued neighboring locations (12.5% each).

Each block consisted of 10 trials of the same cue condition, and each new block started with a different cue condition and the cued locations appeared on the opposite side of the previous cued locations (i.e., the left visual field if the right visual field was cued in the previous block, and vice versa). The order of cue conditions and sides was randomized across blocks and participants via Latin square sequences. Participants were allowed to take short breaks in between blocks without moving their heads from the chin rest. Feedback on performance was given to the participants after every 6th block including the overall accuracy rate and the number of times, they broke fixation since the last feedback. In the main session, participants took a longer break for a few minutes every 18 blocks and they continued the experiment after eyetracker re–calibration. There were 720 trials in total, divided into 72 blocks and by 3 long breaks.

Eye fixation was monitored online with the eyetracker throughout each trial and if participants blinked or broke fixation more than 0.75 dva away from the fixation target during a trial for longer than 0.05 seconds, these trials were marked and discarded in the later analysis. Participants were allowed to move their eyes freely during the response period and ITI.

### Data Analysis

All statistical tests were run using the packages rstatix (Kassambara, 2023) and BayesFactor (Morey et al., 2018) on R v4.3.1 (Team, 2010). To assess how the attentional focus affected discrimination accuracy, a two-way repeated-measures ANOVA with the factors of focus condition (narrow, broad, split) and probe type (cued, uncued) was performed. ANOVA results were Greenhouse-Geisser corrected and reported accordingly with a notation “GG” in the subscript (e.g., GG error as *Sgg)* if the respective term violated sphericity which was determined by Mauchly’s test. As a follow-up, we performed pairwise *t*–tests to determine which conditions drove differences in performance. Furthermore, as we are particularly interested in the differences in performance at the intermediate uncued location compared to the cued and outer–uncued locations we tested these conditions separately. Significance values for the follow­up tests were adjusted through Benjamini–Hochberg procedure denoted as *pBH*. For the non­significant results, we further comparedeach term to a null model that is the same model except for the non-significant term. The deviation from the mean is reported and plotted as ±1 within–subject standard error (*se*; (Morey, 2008)).

The behavioral and EEG data of Experiments 1-3 can be found in the following link: https://osf.io/56er7/?view_only=097fe07e23064a499206660876d89ca4.

### Results

Figure 2 shows the results of Experiment 1. Data from two participants were not included in the final analysis because the participants broke fixation or moved their eyes in more than 25% of the trials. Data from one additional participant was discarded due to the thresholding procedure resulting in a much higher contrast level (59.29%) than that of the rest of the sample (*mean* ± *se*: 12.94% ± 1.25%). For all other participants, average performance was 75% ± 1.89% accuracy as intended by our thresholding procedure. A two-way ANOVA revealed a significant main effect of cue validity (*F*(1,26) = 19.01, *p* = 10^-4^, *η*p^2^ = 0.42, *BF* = 8304) such that participants were overall better at discriminating the cued targets as compared to the uncued targets. Unexpectedly, there was no significant effect of attentional focus condition and no interactions (all *p*’s > 0.2, see Table 1).

**Table 1.** ANOVA Results for Experiment 1.

| | *F* | $\epsilon_{GG}$ | *df* | *p* | $\eta_p^2$ | *BF* | sig. |
| --- | --- | --- | --- | --- | --- | --- | --- |
| **Focus** | 0.59 | 0.968 | (1.94, 50.31) | 0.55 | 0.03 | 0.1 |  |
| **Cue** | 19.01 | | (1, 26) | $10^{-4}$ | 0.42 | 8304 | *** |
| **Focus x Cue** | 1.57 | 0.95 | (1.9, 49.38) | 0.22 | 0.06 | 0.31 |  |

*Note.* The Greenhouse-Geisser sphericity corrections were applied when necessary and the results were reported accordingly.

**Figure 2.**
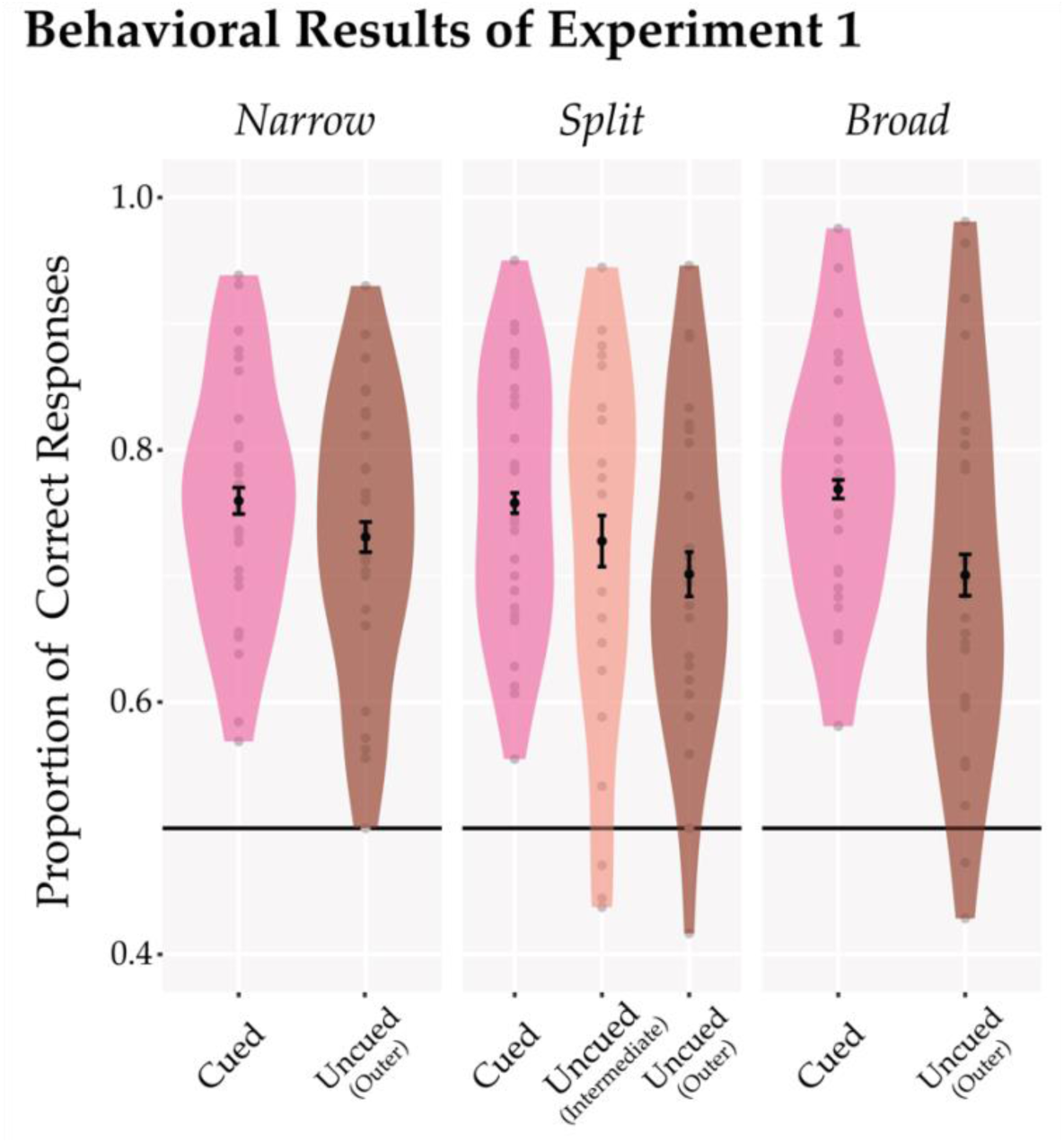
Behavioral results from Experiment 1. The proportion of correct responses (y-axis) is shown separately for each focus condition and cued and uncued locations. Across all conditions, performance was higher for test probes appearing at the cued vs. outer uncued locations. However, the intermediate uncued probes in the split-focus condition (central plot) did not yield different performance levels compared to the cued or the uncued test probes.

Nonetheless, we performed exploratory analyses to examine the effect of cue at each of the attention conditions. The cueing effect in the narrow–focus condition marginally failed to reach significance but there was a trend with a small effect size, *t*(26) = 1.96, *p* = 0.06, *Cohen’s d* = .38, *BF* = 1.06. There was a significant and moderate effect of cue in the broad–focus condition, *t*(26) = 3.77, *pBH* < .004, Cohen’s *d* = 0.73, *BF* = 38.98. When testing the split–focus condition we treated the intermediate and outer uncued locations separately. Pairwise *t*–tests demonstrated that performance at the cued location was significantly higher than at the outer uncued location (*t*(26) = 2.71, *pBH* = .03, Cohen’s *d* = 0.52, *BF* = 4.04). Interestingly the intermediate location did not differ significantly from either location (see Table 2). Additionally, we ran a *t*–test to examine whether the participants performed differently at the intermediate location when it was cued in the broad–focus condition compared to when it was uncued in the split–focus condition. We only included those trials where the probes appeared in the intermediate location. There was a marginally non-significant effect and the trend showed that participants performed slightly better in the broad–focus condition (77.66% ± 0.02%) than they did in the split–focus condition (72.75% ± 0.02%; *t*(26) = 2.03, *pBH* = .13, Cohen’s *d* = 0.39, *BF* = 1.19). Overall, these data indicate that participants were able to effectively attend narrowly and broadly within a hemifield, but when they were asked to split their attention across noncontiguous locations the results were ambiguous (see General Discussion).

**Table 2.**
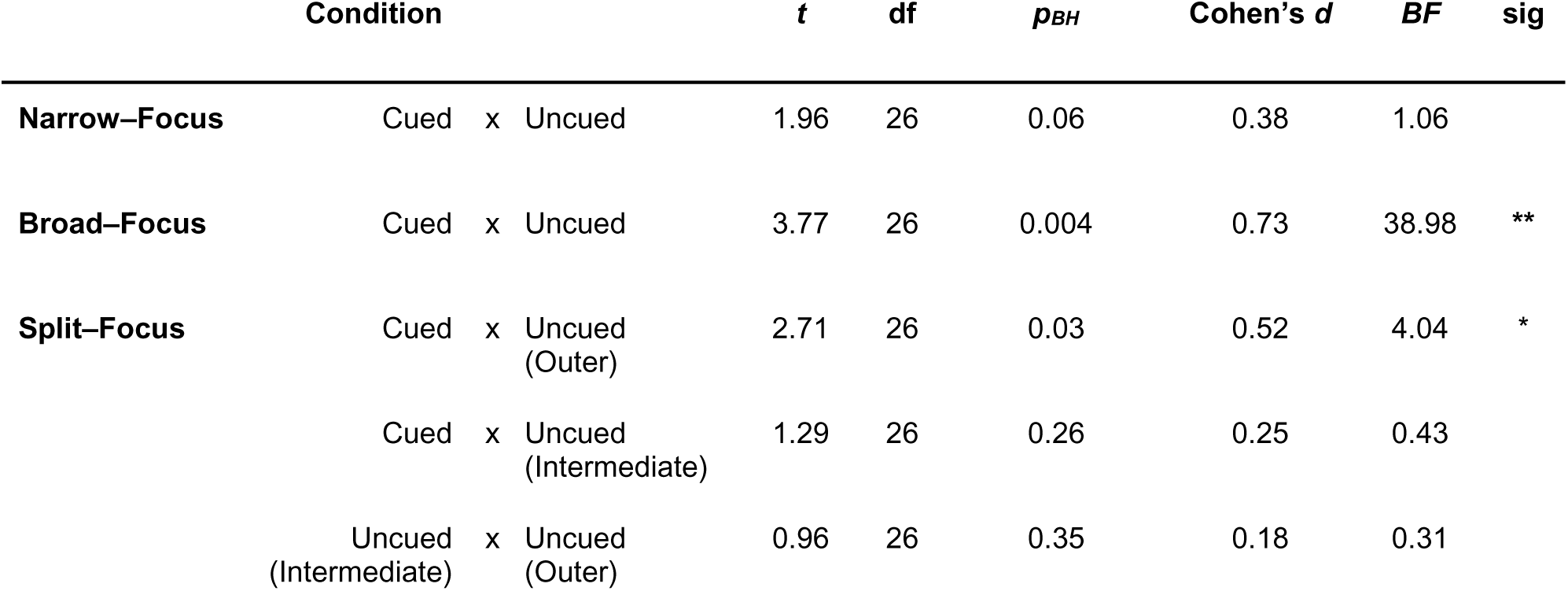
Results from the Post-hoc Pairwise t–tests for Experiment 1.

## Experiment II

In Experiment I, performance at the intermediate to-be-ignored location for the split-focus condition was not reliably reduced, suggesting that attention was not split effectively. It is, yet, possible that attention modulates noncontiguous regions in the early visual cortex, but these differences are obscured at higher levels of processing as assessed with our behavioral paradigm. Thus, in Experiments 2 and 3 we made use of SSVEPs where the attended and unattended locations flickered at distinct frequencies, allowing us to measure the early visual processing of each location directly.

### Methods

#### Participants

Thirty–one undergraduate students from Dartmouth College, aged between 18 and 35 years, participated in the experiment. All participants had normal or corrected-to-normal vision. Just like in Experiment 1, participants signed an informed consent approved by the Institutional Review Board at Dartmouth College prior to the experiment. The protocol was approved by the Committee for the Protection of Human Subjects.

#### Apparatus

In Experiments II and III testing was completed on a PC running Windows 10. The experiment was displayed in an otherwise dark and electrically shielded booth on a ViewPixx monitor (1920 × 1080 pixels at 120 Hz; VPixx Technologies Inc.) placed 38 cm from the participant. The participant’s head was stabilized by a chin rest. EEG was continuously recorded from 32 Ag/AgCl electrodes mounted on an elastic cap, and the signal was amplified by a BrainVision ActiCHamp amplifier (BrainProducts, GmbH). Electrode configuration was based on a 10-20 system except that three frontal electrodes (FC5, FCz, FC6) were placed below the occipital electrodes (as I3, Iz, I4) to increase spatial sampling from the posterior scalp. The horizontal electrooculogram (HEOG) was recorded through a bipolar pair of electrodes positioned next to the lateral ocular canthi and grounded with an additional electrode placed at the right side of the neck. FP1 electrode above the left eye measured vertical electrooculogram (VEOG). The scalp electrodes were referenced online to an electrode on the right mastoid. The recordings were sampled at 500 Hz with an online high–pass filter of 0.01 Hz and low–pass filter of 250 Hz.

#### Stimuli

Stimuli were presented on a salmon-colored background (RGB: [160, 80, 64]). The background color was chosen to make it more pleasant for the participants’ eyes. The layout was created similarly to Experiment 1. Participants were instructed to maintain their gaze on a central fixation point, and the inner circle changed color to indicate the beginning and end of the trial epochs. The lower visual hemifield was divided into 6 equally sized (25.33° of arc) segments separated by 4° from one another and 2° from the vertical and horizontal meridians. The inner and outer edges of the segments extended 6 and 10 dva from the fixation point. A scaled version of the segmented arcs was placed between 2 and 3 dva away from the fixation, serving as a spatial cue: the to-be-attended locations were cued by highlighting the corresponding locations of the inner arcs in yellow (RGB: [181, 179, 92]). As only either the arcs on the left or the arcs on the right visual hemifield were relevant on a given trial, only these arcs were tagged with different frequencies and flickered from white to black at ∼10.91 Hz (innermost), 12 Hz (intermediate), or ∼13.33 Hz (outermost) during the testing period (see *Procedure*). The three different frequencies allowed us to measure the early visual processing separately for each of these locations (see *Analysis*).

#### Procedure

Each session started with the task instructions and a brief (<5 min) practice run of the main task. Then, participants were capped, and electro-gel was applied to each scalp electrode until the impedance was lower than 10 kΩ. When the experiment started, participants were instructed to keep their eyes on the fixation circle and try not to blink throughout each trial. The experimenter monitored the EEG and HEOG recordings and gave participants feedback if they noticed blinks or eye movements during the recording.

As in Experiment I, the study used a block design in which the cues alternated sides (left and right visual half-field) and the focus condition always changed after 5 consecutive trials (480 trials in total) in a random order using Latin square sequencing. At the start of a 5 –trial block, the cue display indicated the to–be–attended locations. The trials were initiated when participants pressed the spacebar and the fixation color turned red as an indication. At the start of the trial there was a 1–sec–long fixation period after which the locations started flickering at their respective frequencies. The trial duration was jittered between 4.5 and 5 secs. Participants were instructed to monitor for a brief dimming of one or two locations (0.4 seconds) at the cued locations. There were 1, 2, or 3 incidences of these dimming events per trial, and only half of them were presented at the cued locations. Participants were instructed to respond as quickly as possible to these target events at the cued locations by pressing the spacebar but withhold any response to nontarget events occurring at the uncued locations. The timing of each event was randomized within the trial with the constraint that any consecutive event onsets were separated at least by 1.1 secs and that no event would occur within the first 0.5 and the last 0.7 secs of a trial. Target and nontarget events varied according to the attention condition: In the narrow –focus condition, either the innermost (with respect to the vertical meridian) or the intermediate location was cued, and a target event was defined as the brief dimming of one of these cued locations; dimming events at any of the uncued locations were nontargets. For the narrow-focus condition, we never cued the outermost location to maximize the number of trials per location/frequency. In the split–focus condition (Figure 3A, left), the innermost and the outermost locations were cued but the intermediate location was uncued, and the target event was the synchronous dimming of the two cued locations. There were two configurations of nontarget events in the split–focus condition. In the first type (singular), only a single location dimmed; in the second type (adjacent), one of the cued locations dimmed along with the intermediate uncued location. Each non–target event was equally probable. In the broad–focus condition (Figure 3A, right), all three locations were cued, and a target event was defined as any two of the three cued locations dimming synchronously and a nontarget event was defined as any single location dimming. Thus, the split– and broad–focus conditions shared one common target configuration (i.e., two *disjoint* locations dimming), and one common non–target configuration (i.e., *singular* location dimming). However, they diverged in one other configuration when *adjacent* locations dimmed together. The adjacent configuration was a target in the broad–focus condition but a nontarget in the split–focus condition. Thus, in order to not false-alarm to the adjacent non–targets in the split-focus condition, it would be beneficial to split attention whereas it would be beneficial to attend to all locations equally to detect both target types (adjacent and disjoint) in the broad–focus condition. The amount of dimming, which was implemented by gradually decreasing (or increasing) the transparency (i.e., alpha value) of the stimulus by following a Gaussian across time (reaching its maximum at the halfway point: *μ* = 0.2 sec, *σ* = 1), was thresholded for each participant throughout the session via a 3-down-1-up procedure (Prins & Kingdom, 2018) for each condition separately. The algorithm was updated after every target event, but the same value was used for the distractor event as well. We also gave participants feedback on their overall performance after every 30 trials in terms of their hit rates, false alarm rate, and the number of unpaired responses given (i.e., no target or distractor event occurred 1 second prior to the response).

**Figure 3.**
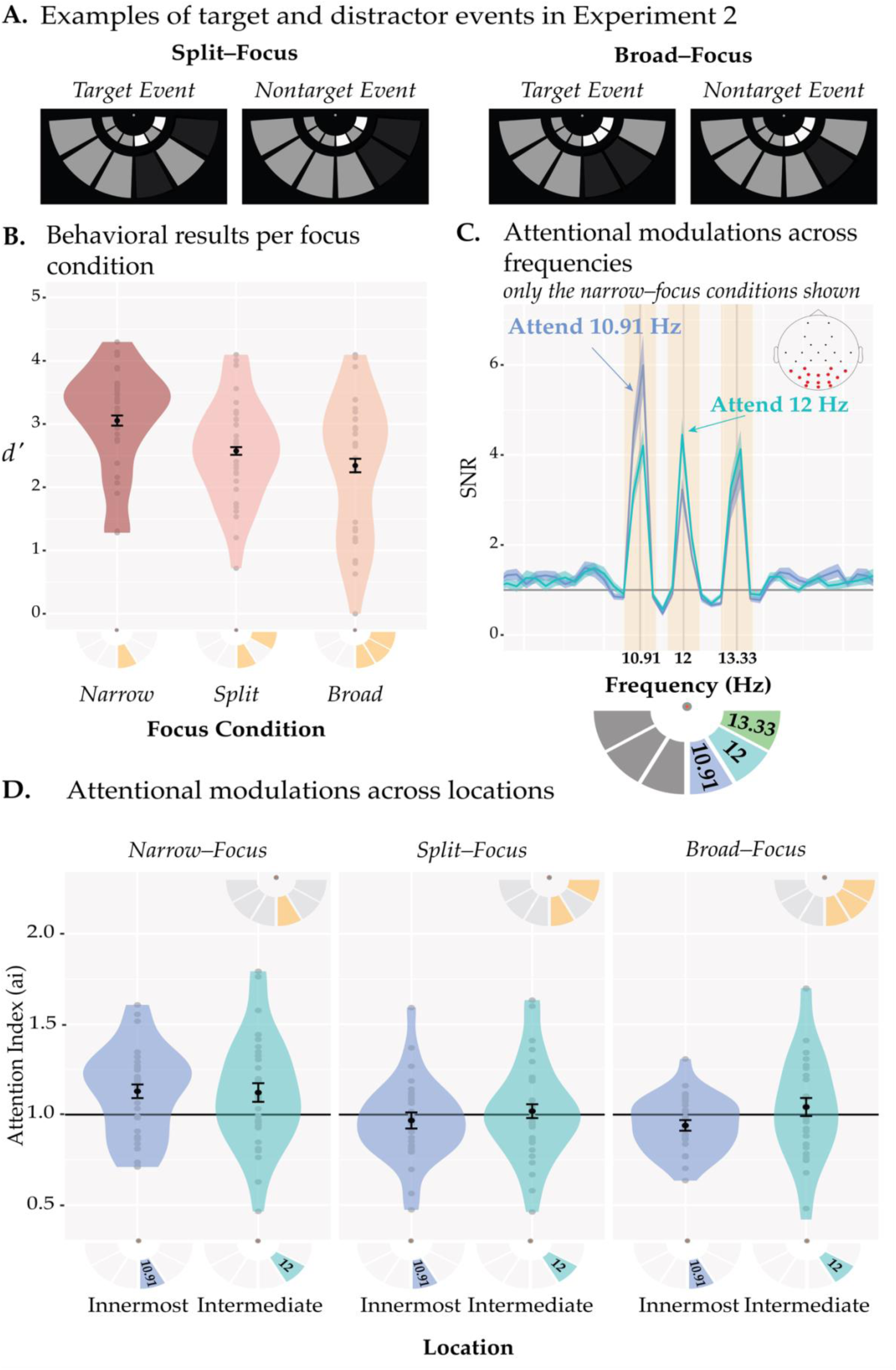
Stimuli and results of Experiment 2. (A) *Examples of target and nontarget events for the split– and broad–focus conditions.* In either event one or more arcs were briefly dimmed. In target events either both (split–focus) or any two (broad–focus) of the cued arc dimmed simultaneously whereas during nontarget events, either a single arc (broad– & split–focus) or the intermediate arc and one of the cued arcs (split–focus) dimmed together. The colors were converted to grayscale for illustration purposes. (B) *Behavioral performance (d’s) across the different focus conditions.* The narrow–focus condition resulted in highest *d’* whereas performance was lower for the split– and broad–focus conditions which did not differ from one another. (C) *SSVEP SNRs in the narrow–focus conditions.* SNRs are shown for two narrow–focus conditions where participants attended the 10.91–Hz stimulus (innermost location) or the 12– Hz stimulus (intermediate location). SNRs at a given frequency were increased when the respective stimulus was attended relative to when it was unattended. The inset on the top right shows the electrode locations, with the electrodes used in the analyses highlighted in red. The search window to determine the peaks for each frequency is denoted by yellow boxes centered around the stimulation frequencies. The stimulation frequencies and their locations are shown in the pictogram under the plot. On half of the trials the order was flipped (left–side task– relevant) along the vertical axis. (D) *Attentional modulations across locations per focus condition.* The baseline attentional modulation (*ai* = 1) is indicated by the black horizontal line. Only the narrow–focus condition induced reliable attention effects, evident as above-baseline SNRs. There was no difference between the two locations (innermost and intermediate). All the error bars indicate ±1 within–subject standard error.

#### Data Analysis

##### Behavioral data analysis

Since participants responded continuously throughout the test interval each response was matched with a target or a nontarget event if it followed the event onset within a 1-sec response window. Only these responses were analyzed further (false alarms that occurred when no dimming was present were discarded from the main analysis). Hit rates and false alarm rates were computed from the responses to target and nontarget events, respectively, and d–primes (*d’*) were calculated and corrected according to the log–linear rule (Hautus, 1995) for each condition separately. A one-way repeated-measures ANOVA with the focus conditions (narrow, broad, split) as a factor was performed. As a follow-up, we performed pairwise t-tests to determine which conditions drove differences in performance. GG–corrections, BH–adjustments, and Bayes’ factor calculations were implemented similarly to Experiment I.

##### EEG data preprocessing and analysis

In the first step of preprocessing, we visually inspected the raw signal from each channel and determined whether any channel contained excessive noise. If so, those channels were marked and made exempt from further analysis. In both experiments, no more than 1 channel was discarded in less than 10% of the participants’ recording. Any marked channels were interpolated via spline interpolation prior to epoching the data. Second, an independent component analysis (ICA) with 29 components (1 minus the number of channels) were implemented to correct for ocular artifacts (i.e., blinks and horizontal eye movements). The resulting independent components and their scalp distribution were inspected visually and those components that resembled blinks and eye movements with activity clustered in the frontal electrodes were excluded (1 or 2 components per participant). Later, the EEG signal was re– referenced to the average signal across all the electrodes, and a low–pass (70 Hz) and a high– pass (.1 Hz) finite impulse response (FIR) filter were applied. EEG data were then epoched from 0.1 sec before to 4 sec after the trial onset and each epoch was baselined (0.1 to 0s with respect to the stimulus onset) and linearly detrended. The same preprocessing steps were implemented in Experiment 3.

For the main analysis, we first averaged all trials separately for each condition and then computed the SSVEPs through a fast Fourier transform (FFT) between the time interval of 0.15 - 4 sec after the flicker onset. To compute the stimulus–induced signal–to–noise ratio (SNR) we first estimated the background activity/noise due to non-stimulus-related processes by applying a median filter (±1 frequency bin or ± 0.24 Hz) to the FFT power (i.e., amplitude raised to the 2nd power). This ensured that the flicker–induced activity would not spill over to neighboring frequencies, and a median filter should eliminate sharp peaks that do not persist across neighboring frequencies. The SNRs were then computed as the ratio of FFT power to the median filtered FFT power. We averaged across 15 occipito-parietal electrodes (I3, Iz, I4, O2, Oz, O1, PO7, PO3, POz, PO4, PO8, P8, P4, P3, P7) for our main SSVEP analysis. These electrodes were selected a priori based on a pilot study using the same frequencies where we implemented a clustering analysis to find a group of electrodes that yield highest SNR per frequency. To determine the peak amplitudes, we used a search window of ± 0.24 Hz around each stimulation frequency and took the maximum value. In order to quantify the attentional modulation for each condition and to allow cross-frequency comparisons, we used the narrow–focus attention condition as a baseline and computed an attention index (*ai*) for each condition by dividing the SNRs of the attended frequency by the mean of the SNRs for that frequency when it was attended and unattended in the narrow–focus condition (e.g., *a/*12^ = SNR(12 Hz | Attend 12 Hz) / mean(SNR(12 Hz | Attend 12Hz, narrow), SNR(12 Hz | Attend 10.91 Hz, narrow))). Thus, the narrow–focus condition always served as the baseline. Since there was no narrow–focus condition where the outermost location (13.33 Hz, see above) was cued, we did not further analyze this frequency. We performed a series of one-tailed *t*–tests to see whether the attention index in each focus condition and location was significantly above baseline. We performed a two– way repeated–measures ANOVA with the factors of focus condition (narrow, split, broad) and location (innermost, intermediate). The rest of the statistical analyses followed the same structure as the behavioral data analysis.

### Results

#### Behavioral Results

Figure 3B shows the behavioral results of Experiment 2. Data from one participant were excluded from the final analysis due to not showing clear peaks at the SSVEP frequencies after the SSVEP data were collapsed across all trials and conditions. A one-way ANOVA with the factor of focus condition revealed that participants’ *d’* varied across different attention conditions (*F*(1.43, 41.6) = 18.37, *pGG* < 10^-4^, *ηp*^2^ = 0.39, *εGG* = 0.717, *BF* = 20294). A series of post-hoc t- tests showed that participants performed better in the narrow–focus condition (*d’* = 3.05 ± 0.08) than in the broad–focus (*d’* = 2.34 ± 0.11, *t*(29) = 4.88, *pBH*< 10^-4^, Cohen’s *d* = 0.89, *BF* = 668.88) and split–focus (*d’* = 2.57 ± 0.06, *t*(29) = 6.28, *pBH* < 10^-5^, Cohen’s *d* = 1.15, *BF* = 23186.12) conditions and the latter two conditions did not significantly differ from each other (*t*(29) = 1.83, Cohen’s *d* = 0.33, *pBH =* .08, *BF* = 0.84). Thus, despite continuous thresholding to adjust performance (the lowest alpha values when averaged across the session: narrow–focus: 44.93 ± 2.54, split–focus: 43.5 ± 1.7, broad–focus: 39.58 ± 2.43; no significant difference, *p*=.8), the narrow–focus condition resulted in higher performance than the other two conditions.

#### EEG Results

Figure 3C plots the SNR of the SSVEPs in the narrow–focus attention condition, illustrating that the stimulation frequencies elicited reliable SSVEPs. When participants attended the innermost location (blue) an increase in SNRs can be observed at the respective frequency (10.91 Hz) as compared to when they attended the intermediate location (turquoise, 12 Hz), and vice versa.

Figure 3D shows the attention indices per location and focus condition. To test whether each focus condition successfully evoked attentional modulations at each location, we first conducted one-tailed *t*-tests. For the narrow–focus condition we found a reliable attention effect at both the innermost (*t*(29) = 2.88, *p* = 0.003) and at the intermediate locations (*t*(29) = 2.18, *p* = 0.02). However, the split and broad conditions did not show reliable attention effects and remained around the baseline level of 1 (all *p*’s > .45). A two–way ANOVA with the focus condition and the location as factors revealed a main effect of the focus condition (*F*(1.82, 52.92) = 12.41, *ηp*^2^ = 0.3, *pGG* < 10^-4^, *εGG* = 0.912, *BF* = 27.09), but there was no main effect of location (*pGG* = 0.38, *BF* = 0.45) and no interaction (*pGG* = .22, *BF* = 0.2). A series of post-hoc *t*–tests (see Table 3) revealed that the narrow–focus condition had a higher attention index (*ai* = 1.13 ± 0.03) relative to the broad–focus (*ai* = 0.99 ± 0.02, *t*(29) = 4.02, *pBH* = 0.001, *BF* = 77.97) and split–focus (*ai* = 0.99 ± 0.02, *t*(29) = 4.01, *pBH* = 0.001, *BF* = 76.73) conditions, and the latter two conditions did not differ significantly from each other (*pBH* = .09, *BF* = 0.2).

**Table 3.** Results from the Post–Hoc Pairwise t–test on Attention Indices in Experiment 2.

| Condition | | | $t$ | df | $p_{BH}$ | Cohen's $d$ | BF | sig |
| --- | --- | --- | --- | --- | --- | --- | --- | --- |
| Narrow | x | Split | 4.01 | 29 | 0.001 | 0.73 | 76.73 | ** |
| Narrow | x | Broad | 4.02 | 29 | 0.001 | 0.73 | 77.97 | ** |
| Split | x | Broad | 0.09 | 29 | 0.93 | 0.02 | 0.2 |  |

**Table 4.** Results from the ANOVA test on Attention Indices in Experiment 3.

|  | *F* | *ε*<sub>GG</sub> | *df* | *p* | *η*<sub>p</sub><sup>2</sup> | *BF* | sig. |
| --- | --- | --- | --- | --- | --- | --- | --- |
| **Focus** | 1.09 |  | (1,23) | 0.31 | 0.04 | 0.27 |  |
| **Location** | 10.7 | 0.937 | (1.87, 43.09) | $< 10^{-3}$ | 0.32 | 9256 | *** |
| **Focus x Location** | 0.38 | 0.926 | (1.85, 42.59) | 0.67 | 0.02 | 0.14 |  |

*Note.* Greenhouse-Geisser sphericity corrections were applied when necessary and the results were reported accordingly.

**Table 5.** Results from the Post–Hoc Pairwise t–tests on Attention Indices in Experiment 3.

| Condition | | | $t$ | df | $p_{BH}$ | Cohen's $d$ | BF | sig |
| --- | --- | --- | --- | --- | --- | --- | --- | --- |
| Intermediate | x | Innermost | 4.04 | 23 | 0.002 | 0.83 | 63.06 | ** |
| Outermost | x | Innermost | 1.08 | 23 | 0.29 | 0.22 | 0.36 |  |
| Outermost | x | Intermediate | 3.35 | 23 | 0.004 | 0.68 | 14.3 | ** |

Together, these results show that there was a reliable attention effect on early visual processing when participants attended to a narrow location, but no such modulation was present for neither broad– nor split–focus of attention. In fact, these two conditions did not differ in their activation profile as indexed by SSVEPs, possibly suggesting that when participants are cued to split attentional resources across two non-contiguous regions, attention acts similarly as if participants are cued to attend broadly across all locations. However, there was some uncertainty on how strongly the task encouraged participants to split attention when cued to do so, given there was no distractor condition that explicitly probed the intermediate location, and thus there might have been only a small cost in performance when participants attended broadly to all locations. Furthermore, the lack of a difference between split and broad attention conditions is difficult to interpret given the lack of attentional modulation with respect to the baseline. This may be due to our choice of the baseline condition: it is unclear whether the mean SSVEP signal of the attended and unattended location in the narrow–focus condition is a valid baseline and might have obscured any attention effects.

## Experiment III

To address the concern of a valid baseline, we ran another SSVEP experiment that focused on the broad– and split–focus conditions only. This time, we flickered all 6 locations at distinct frequencies including the ones on the task–irrelevant side and used the SSVEP responses of these unattended locations to create a more appropriate baseline within each focus condition. Specifically, each location flickered at one of six distinct frequencies eliciting six separable SSVEP signals (see Figure 4A). Across trials, we varied the visual half-field participants were attending to (either broadly or with a split–focus), and we quantified the attentional modulations by dividing the SNR of a particular frequency when that frequency was attended by the mean of the SNRs across attended and unattended trials for that same frequency and focus-condition. Moreover, as there was some concern about how detrimental not-splitting attention would be in Experiment 2, we also changed the task to encourage participants even more to continuously monitor the cued locations by using a rapid serial visual presentation (RSVP) paradigm.

**Figure 4.**
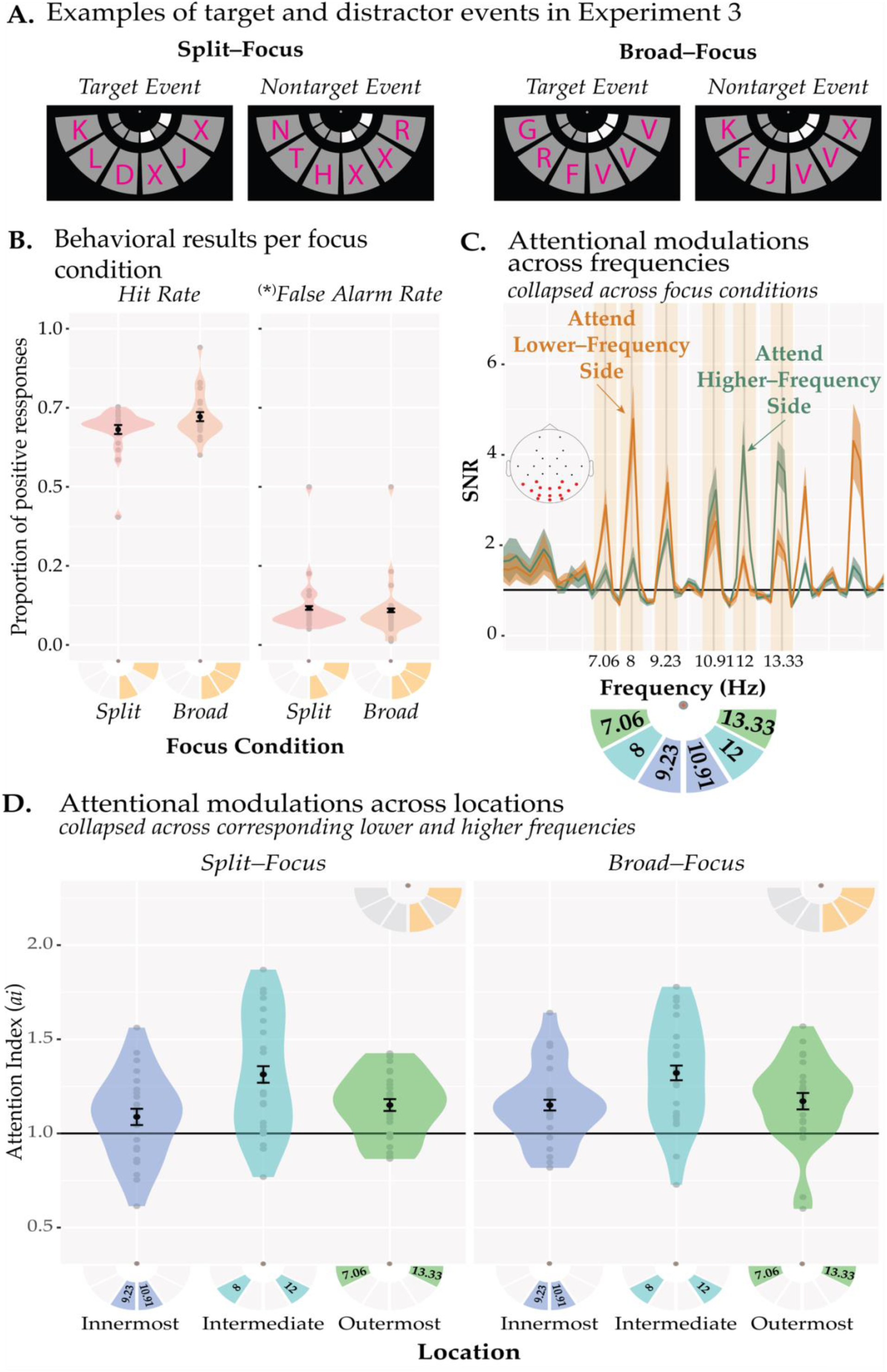
Stimuli and results of Experiment 3. (A) Examples of target and nontarget events. A target event was defined such that all the cued locations had the same letter. We computed false alarms based on the nontarget events where an uncued location and only one of the cued locations contained the same letter (randomly chosen from the 15 letters) in the split–focus condition, and any two of the three locations contained the same letter in the broad –focus condition. (**B**) Hit rates and false alarm rates are shown as a function of the focus condition. There were no significant differences between the focus conditions in either metric. (**C**) SNRs are shown separately for the trials where participants were cued to attend to the locations flickering in lower–frequencies (i.e., 7.06, 8, and 9.33 Hz, orange) or higher–frequencies (10.91, 12, and 13.33 Hz, green), collapsed across the two focus conditions. The stimulation frequencies and their locations are shown in the pictogram under the plot. On half of the trials the order was flipped (higher–frequencies on the left side) along the vertical axis. (**D**) The attention index is shown for each location and focus condition. The data is collapsed across the stimulation frequencies that correspond to the same relative position (collapsed across left and right hemifield). Each location reliably induced attention effects, indexed as above-baseline SNRs (horizontal black line). The focus conditions did not differ from each other at any of the locations including the intermediate location which was cued in the broad –focus condition but uncued in the split–focus condition. All the error bars indicate 1 standard error. *(*) see Behavioral Data Analysis for Experiment 3 to see how the false alarms were calculated*.

### Methods

#### Participants

24 undergraduate students from Dartmouth College, aged between 18 and 35 years, participated in the experiment. We reduced the number of participants relative to Experiment 2 reasoning that using a valid baseline should increase the attention effects overall. As before, they all signed an informed consent form as approved by the Institutional Review Board at Dartmouth College.

#### Stimuli

The overall configuration of stimuli was just like in the previous experiment except the outer arcs were closer in eccentricity: the inner and outer edges subtended 4 and 8 dva from central fixation. Each of the 6 arcs flickered at a distinct frequency, (7.05, 8, 9.23, 10.91, 12, 13.33 Hz in clockwise order from the right-most location on half of trials and in counter–clockwise order from the left–most location on the other half), which allowed us to assess attention effects for each location separately. Participants continuously monitored a rapid serial visual presentation (RSVP) of letters at the attended locations: During the testing period, each of the locations contained a letter randomly assigned from 15 consonants (B, C, D, F, G, J, K, L, N, P, R, T, X, Y, V) in its center. Each letter filled an invisible 1.51 –by–1.35 dva frame. To avoid color adaptation effects and blurring of the stimuli across time, the letters were presented in one of 3 colors (RGB: red [205, 0, 0], dark blue [50, 28, 200], turquoise [0, 180, 155]) randomly assigned at the beginning of each trial. The colors were randomized across all conditions.

#### Procedure

Each block started with the cue display and participants pressed the spacebar to initiate each trial. A trial started with a 1–sec–long fixation period which was indicated by the fixation color turning red. Subsequently, all arcs started flickering and the RSVP stream began. No targets were displayed during the first 0.2 - 0.5 sec (randomly jittered pre–test interval). During the practice and thresholding tasks, the testing period continued for 1 sec where a target letter combination had 50% chance of occurring in each trial. Participants were asked to press the spacebar as soon as possible when they detected a target event. The target event (examples depicted in Figure 4A) was defined based on the focus-condition: for the split–focus condition, the same letter had to be present simultaneously at the two cued locations (e.g., ‘B’ and ‘B’, or ‘N’ and ‘N’ at the cued locations, and any other letter at the intermediate location); for the broad– focus condition, the same letter had to appear simultaneously at all three cued locations (e.g., ‘B’, ‘B’, ‘B’ at all three locations; for a similar design, see McMains & Somers, 2004). Thus, participants were not monitoring for a specific letter, but rather for the co-occurrence of the same letter at all cued locations. We also defined a distractor event for each of the conditions to estimate false alarm rates (also depicted in Figure 4A): For the split–focus condition, a distractor event was defined as any other two locations containing the same letter (e.g., the same letter presented at a cued and uncued (i.e., the intermediate) location); for the broad–focus condition, a distractor event was defined as any two locations containing the same letter (but not all three). Switch rates of the RSVP stream were estimated prior to the main task by updating the thresholding algorithm after every trial that contained a target. We used the nearest possible (depending on frame rate) switch rates acquired in the thresholding procedure except when it coincided with an SSVEP frequency or its harmonics —in which case we adjusted it by ±1 frame (0.0083 Hz). After thresholding, we proceeded with the EEG capping, followed by the main experiment. For the main experiment, the trial structure was similar to that of the thresholding procedure except for the following: the pre–test interval was jittered between 0.5 - 1 sec; the test interval was 4 secs long; there were 0, 1, or 2 target events per trial. The experiment consisted of 480 trials (240 trials per focus condition when left and right sides collapsed). Each block contained 3 trials after which the order of the frequencies was flipped (i.e., clockwise from right–most to counter–clockwise from left–most location, and vice versa). The positions of the frequencies relative to each other were kept constant while every frequency appeared on the left side on half of the trials and on the right side on the other half of the trials. The focus condition was randomized across blocks. Feedback (overall hit and false alarm rate of the trials since the last feedback) was given every 15 trials or 5 blocks.

#### Data Analysis

##### Behavioral Data Analysis

If participants responded within 0.1 to 1 sec after a target or nontarget onset, their response was counted as a hit or false alarm, respectively. We compared hit rates and false alarm rates between the focus conditions separately via a paired *t*–test.

##### EEG Data Analysis

EEG data processing steps were the same as in Experiment 2 except for the way in which the attention index (*ai*) was computed. We first averaged the trials for each frequency and focus condition separately but collapsed across left and right sides. Then, for each frequency and focus condition, the SNR of the attended frequency was divided by the mean of the SNR when attended and the SNR of the same frequency when unattended (i.e., on the uncued side). For example, to compute the *ai* in the high-frequency innermost location (10.91 Hz) for the broad–focus condition (see Figure 4), the SNR at 10.91 Hz in the broad–focus condition when attended was divided by the mean of SNR at attended and unattended-10.91 Hz signal in the broad–focus condition (i.e., *ai*(10.91Hz) = SNR(10.91 Hz | Attended 10.91 Hz) / mean(SNR(10.91 Hz | Attended 10.91Hz), (SNR (10.91 Hz | Unattended 10.91-Hz)). After the *ai* was computed for each frequency and focus–condition, we collapsed these *ai*s across the corresponding locations: 9.23 and 10.91 Hz for innermost, 8 and 12 Hz for intermediate, 7.06 and 13.33 for outermost (see the inset in Figure 4C).

### Results

#### Behavioral Results

Figure 4B depicts the behavioral results. We found that neither the hit rates (*t*(23) = 2.02, *pBH* = 0.11, *BF* = 1.21) nor the false alarm rates (*t*(23) = 0.92, *pBH* = 0.37, *BF* = 0.31) were significantly different between the broad– and the split–focus conditions. The first five participants did not go through the thresholding procedure and the RSVP switch rate was fixed to 2.5 Hz. The remaining 19 participants were thresholded per focus condition; to reach the same accuracy level in terms of hits and false alarms, the letters had to switch slower during the split–focus relative to the broad–focus condition (2.25 ± 0.09 Hz in the split–focus; 2.88 ± 0.09 Hz in the broad–focus condition; *t*(18) = 4.73, *p* < 10^-3^).

#### EEG Results

Figure 4C depicts the SSVEP responses when participants attended locations on the high–frequency side or low–frequency side —collapsing across both focus conditions. We found reliable SSVEPs at all frequencies and their amplitudes were modulated by attention, such that when participants attended to locations in the visual field that was driven by the higher-frequency flickers (orange), SNRs were enhanced relative to lower-frequencies (green), and vice versa.

We, first, tested whether the SSVEPs were modulated reliably by attention for each of the locations. A series of one–tailed *t*–tests showed that the *ai* at each location and focus condition was above baseline (all *pBH*’s < 10^-15^, Figure 4D), indicating that the early visual-cortical responses were modulated by attention reliably. To test whether the magnitude of the attentional modulations differed across locations and/or focus conditions we ran a two-way repeated measures ANOVA. There was a significant main effect of location (*F*(1.87, 43.09) = 10.07, *pGG* < 10^-3^, *BF* = 9256), but no significant differences between the focus conditions (*p* = 0.31, *BF* = 0.27) and no interaction (*pGG* = 0.67, *BF* = 0.14). Thus, participants appeared to distribute their attention the same way during the broad–focus and split–focus condition. To examine how attention effects differed across the locations, we next collapsed across the focus conditions and conducted post-hoc *t*-tests which revealed that the intermediate location (1.32 ± 0.04 *ai*) showed the largest attention effect relative to both the innermost (1.12 ± 0.03 *ai*, *t*(23) = 4.04, *pBH* = 0.002, *BF =* 63.06) and the outermost (1.16 ± 0.03 *ai*, *t*(23) = 3.35, *pBH* = 0.004, *BF* = 14.3) locations. The outermost and the innermost locations did not differ significantly from one another (*pBH* = 0.29, *BF* = 0.36).

Together, these data indicate that participants were distributing their resources surprisingly similarly in the broad–focus relative to the split–focus attention condition, with strongest attentional modulations at the central intermediate location that was cued in one case (broad) but uncued and to-be-ignored in the other case (split).

## Discussion

The current study investigated how attentional allocation differs when participants are instructed to split resources across two non-contiguous locations or are asked to attend broadly to one large contiguous region within a visual quadrant. We found that attentional enhancement did not differ between the split–focus and broad–focus conditions, suggesting that when participants are instructed to divide their attentional focus across two locations within a visual quadrant, attention acts to enhance the cued but also intermediate uncued locations, just like when attending to a broader region in space. This was true in terms of behavioral performance where we found that the visual discrimination sensitivity did not differ at the intermediate uncued location compared to the cued locations (Experiment 1), as well as in terms of visual-cortical modulations measured with SSVEPs, where we found similar attention effects across these two attentional focus conditions (Experiments 2 & 3). Of particular interest, the profile of attentional enhancement appeared strikingly similar across these conditions such that the intermediate location – which corresponded to the to-be-ignored location in the split–focus condition but to the centrally attended location in the broad–focus condition – showed a higher attention index than its neighboring locations.

### Behavioral performance indicates no effective splitting of spatial attention

The behavioral results from Experiment 1 suggest that participants were not able to divide their focus of attention effectively as they processed the intermediate uncued location in the split– focus condition not reliably differently from the cued locations. This is broadly consistent with previous behavioral studies that examined conditions of divided attention within a single hemifield (Alvarez & Cavanagh, 2005; Awh & Pashler, 2000; Bahcall & Kowler, 1999; Liu et al., 2009; Sereno & Kosslyn, 1991) or visual quadrant (Carlson et al., 2007). For example, when participants were instructed to attend to locations in different visual quadrants and target probes were presented either at these cued locations or at intermediate, uncued locations, participants appeared to not be able to effectively divide their focus of attention, especially within a single visual hemifield (Awh & Pashler, 2000). Even more dramatic effects have been observed in attentional tracking tasks where the capacity to keep track of multiple moving targets amongst distractors appears to be halved when these targets are distributed within relative to across hemifields (Alvarez & Cavanagh, 2005). Broadly, the difficulty in splitting attention within a hemifield is consistent with a model of visual attention where target locations are prioritized independently in each visual half-field, and thus compete more strongly within a hemifield relative to across-hemifields. Such hemifield independence may arise either because of independent attentional pointers in each hemisphere (Alvarez & Cavanagh, 2005; Franconeri et al., 2013), or because of local competition of early spatial representations that are stronger within versus across hemifields (Störmer et al., 2014). Interestingly, in our experiment, behavioral performance at the intermediate location reached an accuracy level in between the performance level of the cued and outer-uncued locations, and in fact did not differ reliably from either one of them. It could be the case that this intermediate performance level at the uncued location reflects a mixture of trials where sometimes resources were split effectively and other times not. It could also be the case that such variability arose at the participant-level, with some participants being able to split their focus effectively and others not. Such variability might be due to several factors such as individual differences in attentional capacity, sampling rate of attention, or the strategy used. Future studies could investigate such trial-by-trial variation or between–subjects variability in more detail.

Surprisingly, we did not observe a difference in the attention effect (cued vs. uncued locations) between the narrow–focus and the other two conditions. Based on the zoom-lens model of attention (Eriksen & St James, 1986), we would have predicted that the strength of visual attention wanes as the attended region increases, and that attentional selection is most effective for a smaller region (Castiello & Umiltà, 1990). What could explain this null result? One possibility is that the stimulus size of the target (the Gabor patch) that we used was not optimal to induce a narrow focus of attention the way we intended. Specifically, the Gabor patch filled the prescribed area entirely and no flankers were included to force participants to maintain a particularly narrow focus. Thus, it could be the case that attentional allocation “spilled over” to the uncued regions and was not as narrowly focused as indicated by the cue, which would result in overall weaker cueing effects in the narrow-focus condition. The fact that the effect size of cueing in the narrow– focus condition was smaller compared to the other two conditions somewhat supports this interpretation.

### No evidence for split attention effects in early visual-cortical processing

Experiments 2 & 3 provided insights into how multifocal attention operates in the early visual cortex through inspection of SSVEPs that allowed the tracking of separate visual responses to attended and unattended locations. To compare effects across frequencies, we computed attention indices for each location/frequency combination by baselining SNRs to an unattended condition for every respective frequency. For the analysis in Experiment 2, we chose the unattended locations during the narrow–focus condition as a baseline to compute the attention effects for all conditions, which revealed reliable attention effects for the narrow–focus, but not for the split–focus nor for the broad–focus conditions. These results might suggest that early visual– cortical processing was not modulated reliably in these conditions (cf., Toffanin et al., 2009), and that they resulted in an overall similarly weak activation strength; however, the experiment left open the possibility that differences would be present but were obscured by the baseline we chose to compute the attention indices. In a similar study, Walter, Quigley, and Müller (2014) used a condition where participants attended a central location as their baseline —which is in some ways similar to the narrow–focus condition in our design. Their experiment also failed to induce reliable attentional modulations when participants attended two locations within a hemifield. Hence, to address this concern, we ran another SSVEP experiment focusing on the comparison between split– and broad–focus only but introducing a more straightforward baseline that compared processing of attended and unattended locations within instead of across the attentional-focus conditions. While we observed reliable attentional modulations across both conditions in this third experiment, the broad– and split–focus conditions resulted in essentially identical attention profiles with the activation strengths peaking at the intermediate locations and dropping at the outer locations — suggesting that participants deployed a unitary focus centered around the intermediate location even when instructed not to do so.

The pattern of neural enhancement for the split-focus condition, where we found the strongest SSVEP response at the uncued intermediate location, diverges from the behavioral results of Experiment 1 which showed intermediate performance for this location (not reliably differing from the cued or outer uncued locations, see Figure 2). There are several reasons that may explain these differences. First and foremost, the SSVEP data assess changes in early visual processing due to attention, whereas any behavioral response is the outcome of several processing stages including early visual processes as well as decision-making and preparing a motor response. As has been argued before, it may well be the case that later processing stages are less affected (or differentially affected) by constraints on early visual processing (Störmer, Alvarez, Cavanagh, 2014; Shioiri et al., 2016). Consistent with this, in Experiment 3, slower letter switch rates were necessary in the split–focus condition to achieve a similar performance level to that in the broad–focus condition. This aligns well with studies demonstrating that during a multiple–object tracking task slower speeds are necessary when tracking targets within a hemifield compared to across hemifields to reach the same level of accuracy (cf. Störmer et al., 2014). Such differences in task difficulty are expected if attention inadvertently enhances irrelevant information (i.e., the intermediate location in the split–focus condition) early on in the visual processing stream, as indicated by our SSVEP results, but later-stage processes – possibly operating at a slower rate – can successfully disambiguate the relevant from the irrelevant information; this can then lead to overall increased processing time (e.g. slower speeds or switch rates as in Experiment 3). Thus, differences between SSVEP responses (Experiments 2 and 3) and behavioral performance (Experiment 1) are likely the result of behavioral measures capturing the accumulation of multiple processing stages, whereas the SSVEP responses are sensitive to early visual-cortical limits.

In addition to these theoretical considerations, there are also several differences between the behavioral task used in Experiment 1 and the tasks used in the SSVEP paradigms (Experiments 2 and 3), such as timing (relatively short trials in Experiment 1 compared to long, sustained periods of splitting attention in Experiments 2 and 3), target and distractor event types and detection vs. discrimination tasks (orientation task in Experiment 1, but relatively complex detection tasks in Experiments 2 and 3). Hence, the neural effects of Experiments 2 and 3 do not necessarily directly map onto behavioral measures in Experiment 1. Most broadly though, they align with our main conclusion that splitting attention within a visual quadrant is not particularly effective.

Overall, the absence of differences across the split– and broad–focus conditions in either of the experiments makes a compelling case that spatial attention operates the same way in both conditions even when it is detrimental to task performance. Broadly, these results support models of attention that assume no effective splitting within a single hemifield (Störmer et al., 2014; Walter et al., 2014), but expand these models by demonstrating that cueing to divide attention results in the same attentional profile in early visual cortex as instructing participants to broaden the attentional focus.

### Mechanisms of allocating attention across different spatial regions

What attentional operations underlie the SSVEP modulations and behavioral performance we observe in our tasks? The similarities between split–focus and broad–focus conditions across our experiments support a model in which attention is allocated broadly in both cases. However, such a model would also assume that attentional enhancement is spatially stationary and temporally sustained. However, there is a possibility that non–contiguous splitting can occur briefly but is not sustainable for several seconds of a trial (Dubois et al., 2009). Consistent with this, Itthipuripat, Garcia, & Serences (2013) tested multifocal attention within a visual quadrant using SSVEPs and did not find any reliable attentional modulations when looking at the signal across the trial. However, when examining the response–locked activity in the time-frequency domain, attention effects emerged shortly before the correct responses only, such that target-related SSVEP amplitudes were higher than the distractor-related ones. The authors’ explanation was that attention was not divided continuously but possibly at brief moments in time, and that if splitting occurred prior to a target event, it was more likely that participants gave a correct response. This type of splitting for short periods in time would be missed when the signal is transformed to the frequency domain across the entire trial, as done here. Unfortunately, as we chose frequencies nearby each other to tag the different locations, we do not have the frequency resolution necessary to examine how the SSVEP signals changed across time as they would not separate clearly in a time-frequency analysis.

Another possibility is that during the split–focus condition, instead of continuously broadening the attentional focus (with possibly brief moments of splitting, as described above), participants attended serially to the three locations with a narrow focus and moved their attentional focus back and forth between all locations. This type of “scanning” behavior could yield similar results as observed in the present study, but it would require that this scanning occurs at a particular rate with a relatively stronger modulation than a broad focus in order to achieve the same pattern as observed during the broad–focus condition. Furthermore, and most critical to the question of interest in the present study, this type of scanning mechanism would have to include the intermediate location to allow for this location to result in the strongest attentional enhancement, as we see in our data; this, again, would speak against an account of effective splitting. In either case, our results show that whatever mechanism supported attentional allocation across the split– and broad–focus conditions, it resulted in the exact same pattern of enhancement across locations in early visual processing.

One potentially interesting question for future research is to explore whether effective splitting could arise through incidental learning processes. Recent research has shown that distractors are often effectively ignored based on statistical regularities, such that over time, locations that contain a frequent distractor are better ignored than infrequent distractor locations (e.g., Chang & Egeth, 2019; Sawaki & Luck, 2013; van Moorselaar & Slagter, 2019; Wang & Theeuwes, 2019). None of these tasks have looked at conditions in which distractors appear between two targets (possibly requiring splitting). While our design does not allow the investigation of such learned attention effects, in future work it could be interesting to test whether incidental learning about relevant or irrelevant locations would result in different, potentially more effective (Wang & Theeuwes, 2019; Addleman & Störmer, 2023), splitting of attention, relative to cued, voluntary attention as assessed here.

## Conclusion

In a series of experiments, we examined the differences in visual discrimination sensitivity and early visual–cortical processing of sustained spatial attention during split and broad cueing conditions within a visual quadrant. Our experiments used different task designs and measurements and yielded consistent findings indicating that resources cannot be split effectively across non-contiguous locations but are instead distributed broadly across relevant and irrelevant locations. Critically, by measuring attentional modulations with high spatial resolution at different locations using SSVEPs, we were able to show that the profile of attentional enhancement is strikingly similar when participants are instructed to divide or broaden their focus of attention, revealing a gradient profile with an activity peak at the central region and a gradual fall-off of this enhancement for nearby attended locations. Our results suggest that sustained spatial attention operates in a unitary focus within a visual quadrant and cannot be effectively divided. Most broadly, our results indicate that the allocation of attentional resources is constrained by the structure and organization of the visual system.

## Declaration of Competing Interests

The authors disclose no conflicts of interest related to this manuscript.

## Author Contributions

Both authors took role in literature review, hypothesis formation, designing the experiments, writing, and editing of the manuscript. The first author programmed the experimental and analysis scripts and collected data.

## Acknowledgments

The authors would like to thank the research assistants Maya Resnick and Justin A. Santana for their help with data collection. We also would like to thank Alireza Soltani for providing access to their eyetracking equipment.

## Supplemental Materials

### HEOG Activity in Experiments 2 and 3

**Figure S1.**
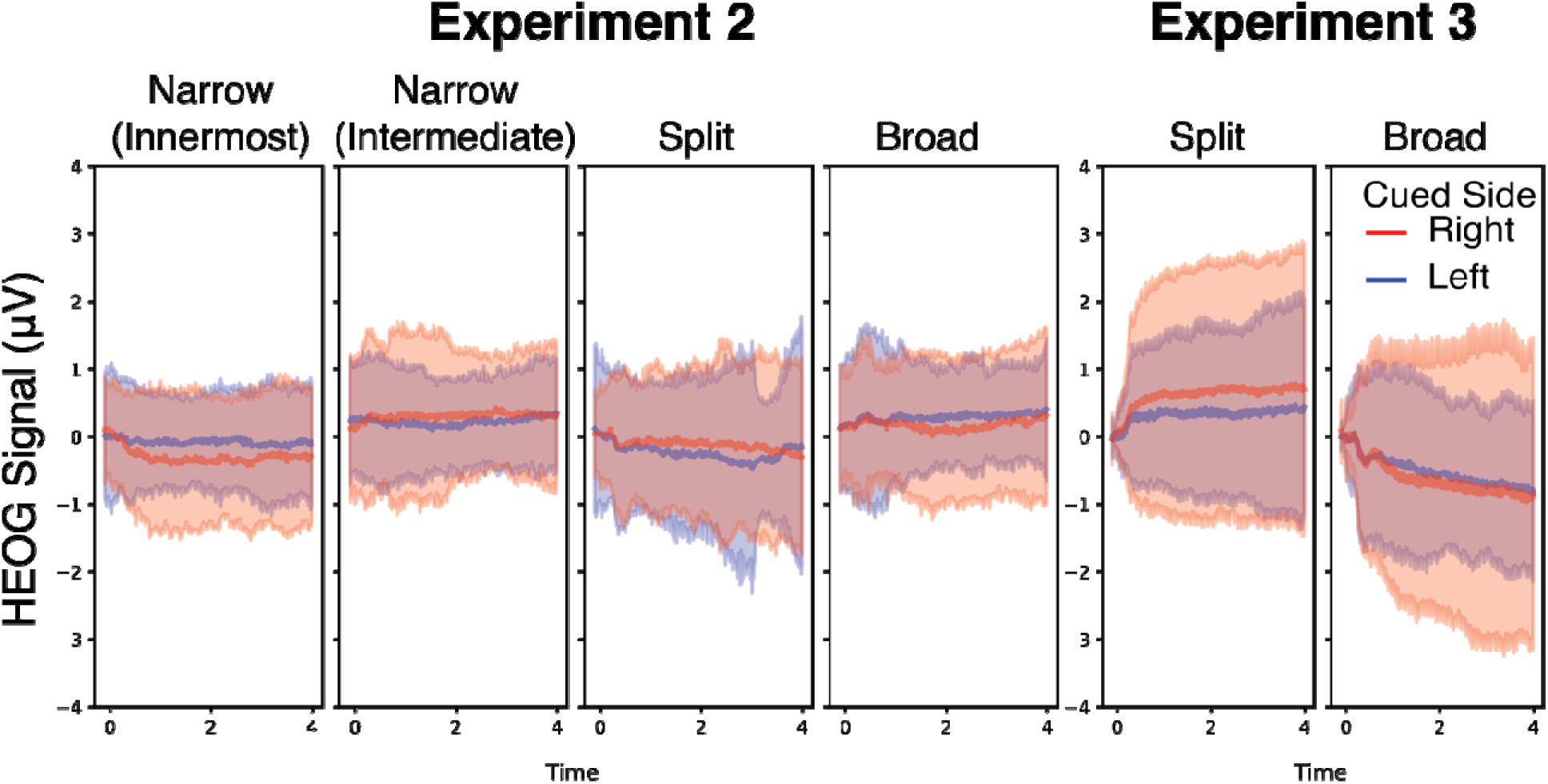
HEOG activity per focus condition and cue side in Experiments 2 & 3. The HEOG signal was averaged across trials per focus condition and the cued side for each participant separately. The shaded areas denote for 95% confidence interval after Morey-Cosusineau correction. Each column represent a focus condition. The narrow-focus condition in Experiment 2 contained two configurations in which either the innermost or the intermediate locations were cued, and these configurations were plotted and analyzed separately. Cue side is color coded such that red represents right-side-cued blocks, and the blue represent left-side-cued blocks. The HEOG signal did not vary with focus condition in either experiment. There was a main effect of cue side in both experiments but see the supplementary text for interpretation.

Here, we analyzed the HEOG activity to test whether participants’ retinal positions varied across different focus conditions and cue side (left or right visual hemifield). HEOG signals were first baselined to the average signal at the interval from -0.1 s until the flicker onset for each trial. Then, we averaged those trials that belonged to the same cue side and focus condition for each participant. Data from 6 participants in Experiment 2 and 5 participants in Experiment 3 were discarded due to faulty signal: the HEOG electrodes were detached from skin during the session yielding unusable data. In Figure S1, we show the group average of HEOG activity across time for Experiments 2 and 3 separately for each focus condition and cue side. Positive values indicate eye movements toward the rightwards direction. To test if there was any significant eye movement depending on the focus condition and the cue side, we averaged the HEOG signal across time and ran a two-way repeated-measures ANOVA. In both experiments we found a significant main effect of cue side (see Table S1 & S2). However, there was no main effect of focus condition or an interaction. In Experiment 2, participants’ HEOG activity was more positive in the right-side cued trials (0.45 ± 0.14 μV) compared to the left-side cued trials (-0.22 ± 0.14 μV, *p_BH_* = 0.009) indicating eye movements towards the cue side. In Experiment 3, participants’ HEOG activity was more negative in the right-side cued trials (-0.56 ± 0.14 μV) compared to the left-side cued trials (0.46 ± 0.14 μV, *p_BH_* = 0.0001) indicating eye movements towards the opposite side of the cue. Both results suggest that the retinal position of the stimulus did not vary depending on the focus condition in either experiment but might have depended on the cue side. However, we should be cautious of interpreting these effects’ significant since the magnitudes of HEOG signal are miniscule and might not correspond to any real eye-movements. According to previous reports a horizontal eye movement of 1-degree visual arc results in ∼16 μV activity^1^. Moreover, the direction of the eye movement was towards the cued side in Experiment 2, but it was opposite of the cued side in Experiment 3, which further reduces interpretability of this result. We, also, ran the main ANOVA tests on attention indices in each experiment excluding the participants with faulty HEOG reading (see Tables S3 & S4). The significance and the direction of the factors remained the same as those in the main analysis pipeline.

**Table S1.** ANOVA results for average HEOG activity per focus condition and cued side in Experiment 2.

| | *F* | $\epsilon_{GG}$ | *df* | $P_{BH}$ | $\eta_p^2$ | *BF* | sig. |
| --- | --- | --- | --- | --- | --- | --- | --- |
| **Focus** | 1.87 | 0.376 | (1.13, 23.67) | 0.18 | 0.08 | 0.13 |  |
| **Side** | 11.36 |  | (1, 21) | 0.009 | 0.35 | 51007510 | ** |
| **Focus x Side** | 1.98 | 0.468 | (1.4, 29.46) | 0.18 | 0.09 | 0.13 |  |

**Table S2.** ANOVA results for average HEOG activity per focus condition and cued side in Experiment 3.

| | *F* | $\epsilon_{GG}$ | *df* | $P_{BH}$ | $\eta_p^2$ | *BF* | sig. |
| --- | --- | --- | --- | --- | --- | --- | --- |
| **Focus** | 0.71 |  | (1,18) | 0.41 | 0.04 | 0.27 |  |
| **Side** | 28.12 | | (1, 18) | $10^{-4}$ | 0.61 | 7992648 | *** |
| **Focus x Side** | 1.75 |  | (1,18) | 0.2 | 0.09 | 0.59 |  |

**Table S3.**
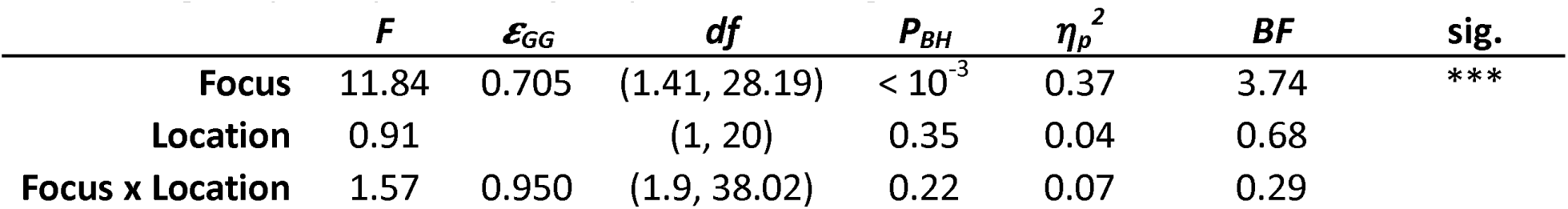
ANOVA results for attention indices per focus condition and cued side in Experiment 2 excluding the participants with faulty HEOG reading.

**Table S4.** ANOVA results for attention indices per focus condition and cued side in Experiment 3 excluding the participants with faulty HEOG reading.

| | *F* | $\epsilon_{GG}$ | *df* | $P_{BH}$ | $\eta_p^2$ | *BF* | sig. |
| --- | --- | --- | --- | --- | --- | --- | --- |
| **Focus** | 1.03 |  | (1,18) | 0.32 | 0.04 | 0.34 |  |
| **Location** | 6.13 | 0.998 | (2, 35.94) | 0.005 | 0.25 | 44.9 | ** |
| **Focus x Location** | 0.13 | 0.862 | (1.72, 31.04) | 0.85 | 0.01 | 0.14 |  |

### SNR Topologies per Frequency

**Figure S2.**
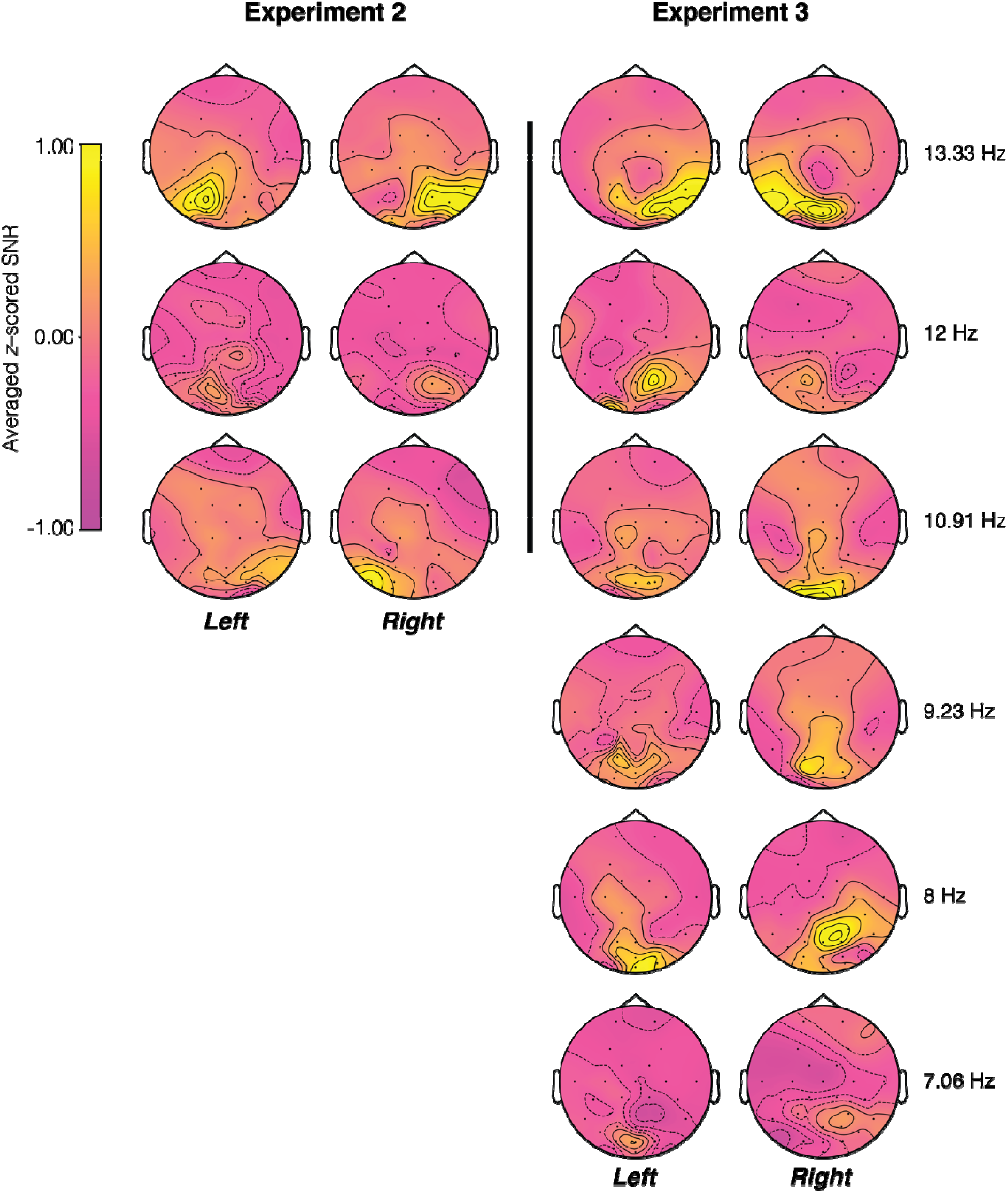
Average SNR topologies for each stimulation frequency per stimulius side. SNR topologies are shown for Experiments 2 (left) and 3 (right) for each presentation side and stimulation frequency, separately. Every participants’ SNRs at each stimulation side and frequency (averaging across focus conditions) were calculated in the same way as in the main analysis. The resulting SNRs were then z-scored across electrodes after which the grand average is taken across participants and plotted here. Stimulation frequencies induced increased signal in the posterior electrodes in both experiments. No consistent lateralization was observed.

## Footnotes

1 page 199 in Luck, S. J. (2014). An introduction to the event-related potential technique. MIT press.

## Notes

https://osf.io/56er7/?view_only=097fe07e23064a499206660876d89ca4

### Competing Interest Statement

The authors have declared no competing interest.

